# Long-lived coherences for magnetic interactions in proteins

**DOI:** 10.1101/2023.07.22.550138

**Authors:** Florin Teleanu, Andrei Ciumeica, Octavian Ianc, Adonis Lupulescu, Aude Sadet, Paul R. Vasos

**Affiliations:** Biophysics and Biomedical Applications Group and Laboratory, ELI-NP, LGED, “Horia Hulubei” National Institute for Physics and Nuclear Engineering IFIN-HH, 30 Reactorului Street, 077125 Bucharest-Măgurele, Romania; Interdisciplinary School of Doctoral Studies (ISDS), University of Bucharest, 36-46 Mihail Kogălniceanu Bd, Bucharest, 050107, Romania and Faculty of Physics, University of Bucharest, 405 Atomiştilor Street, 050663 Măgurele, Romania

**Author notes:** Paul R. Vasos, Aude Sadet. **Email:**.

## Abstract

Living systems rely on molecular building blocks of low symmetry such as amino-acids and nucleotides, which generally yield short-lived magnetic transitions in response to electromagnetic radiation. This is the first demonstration that in proteins of relatively large size local magnetic symmetry can be induced to enable the detection of interactions based on long-lived coherent transitions of nuclear spins. Long-lived coherences (LLC’s) are superpositions of quantum states with singlet and triplet spin-permutation symmetries that feature significantly longer relaxation time constants compared to those of standard nuclear spin coherences. We report in this study that glycine residues in Lysozyme, a 14.3 kDa protein, feature Gly-H^α2,3^ LLC’s with relaxation time constants twice as long as the classical counterparts. Using a new excitation method for LLC’s in glycines 4, 49, 54, 67, 117, and 126, Lysozyme Gly-H^α^ dipolar interactions with neighboring hydrogen spins were mapped in a high magnetic field – at 950 MHz ^1^H Larmor frequency. As predicted by theory, the positions of nearby atoms in the protein structure on one side or the other of Gly molecular symmetry planes reflecting protons H^α2,3^ determines the signs of LLC magnetic interaction signals. LLC-based transfers therefore yield stereospecific signals from glycine residues to ^1^H neighboring atoms. The symmetry-encoded sign of the detected signals provides angle constraints, in addition to the distance information. LLC probes based on naturally-abundant ^1^H spins can be useful for in-cell spectroscopy, circumventing the introduction of heterogenous spin labels for following protein-ligand or protein-protein interactions in the natural environment.

This is the first demonstration that magnetisation transfer through space from long-lived coherences can be obtained in proteins. Applications of LLC’s were believed to be limited to systems featuring fast rotational motion in solution, mainly small molecules. This new LLC-based method yields stereospecific distance constraints and has the potential to extend the protein-size domain for the study of intra- and intermolecular interactions.

## Introduction

The constituents of living matter evolved from primary elements via a process that lowered structural symmetry to enable specific key-lock interactions. Unlike basic oligo-atomic molecules such as H_2_, O_2_, N_2_, CO_2_, few amino-acids feature rotational symmetry elements, and such symmetry is lost when amino-acids are included in a protein fold. The time scale of observable magnetic dipole transitions between symmetry-allowed states in the case of magnetic-dipole emissions for H_2_ is of the order of hours (1). The relationship between structure symmetry and observable timescales of transitions is given by mathematics laws, starting with Emmy Noether’s theorem (2). It was discovered recently that magnetic resonance experiments can be conducted under conditions that establish local symmetry at the observed site, obtaining long-lived spin populations based on singlet states (3) or transitions between permutation-symmetric and antisymmetric states (4). Introducing a magnetic field component to eclipse differences between local magnetic environments in aliphatic protons, as demonstrated in (5), we have shown that local magnetic symmetry can be enhanced for high-field transitions based on singlet/triplet configurations of nuclear spins, namely long-lived coherences (LLC’s) (6). This opened the way for recording ^1^H magnetic resonance transitions with relaxation time constants extended by up to a factor 9, even in peptides or proteins, namely for Gly protons. However, lifetime enhancements for magnetic transitions were only detected experimentally for small molecules and there remained the challenge of demonstrating that such long-lived coherences can be used for magnetization transfer in a protein, though theoretically this has been deemed possible (7). We demonstrate herein that 1H-based LLC’s with lifetimes twice as long as standard coherences, *T*_LLC_ > 2*T*_2_, can be observed in 14 kDa protein Lysozyme and used to transfer magnetisation to remote amino-acids within the protein fold, thereby establishing distance constraints. The intensity of the LLC-based transfer signal is up to 2 times higher compared to the signals obtained by classical transverse magnetisation transfer (8).

This is an important step forward in probing protein structure and protein interactions based on endogenous magnetic probes that can be detected non-invasively. Protein fold and interactions in cells are essential for cell homeostasis and for cell function. The long lifetimes afforded by phosphorescent probes, which can be used for medical imaging, are also sensitive to inter-molecular interactions (9,10). However, most chromophores are difficult to introduce in molecular folds without perturbing the structure. Atomic resolution in liquid state can only be solved non-invasively by Nuclear Magnetic Resonance (NMR). Hydrogens receive deserved credit as first sensors of intramolecular and inter-molecular interactions. Glycine residues are often placed in protein loops, and therefore Gly-based magnetic interactions are of critical importance to define labile structural regions that are difficult to model correctly using AlphaFold (11) or based on solid-state constraints derived from X-ray data. The structural flexibility of glycine-rich protein regions in solution enables biological interactions; many partially-disordered proteins are known to adopt biologically-active conformations relying on their flexibility (12–14). In high magnetic fields, 1H NMR is particularly adapted to follow protein interactions with ligands. Protein resonance assignment, beyond solving the structure, allows interactions to be sensed in pharmaceutical NMR at protein active site. The solution structures of proteins can be solved by magnetic resonance spectroscopy in liquid state with fair resolution, up to a certain size. Biomolecular structure in liquid state beyond proteins sizes of 40 kDa remain elusive (10,15). Efforts in surpassing the detection limit for increasing protein size included the transverse relaxation optimized spectroscopy experiment (TROSY), relying on nuclear transitions in coupled spins of two different nuclei, ^1^H and ^15^N, that feature spin states with long lifetimes (16). The lifetime enhancement in the TROSY experiment is due to the protection of involved nuclear states by the action of relaxation mechanisms -dipole-dipole and chemical shift anisotropy relaxation- of similar strength and opposite effects. Observations of nuclear magnetic transitions in the present study of interactions rely on detecting pairs of ^1^H spins with transitions protected from the spin-spin dipolar interaction (3). Singlet and triplet states are relevant for coupled ^1^H spins in high magnetic field provided a sustaining radio-frequency field of large-amplitude – compared to the difference between local magnetic field of the two protons - is introduced during the experiment. The singlet/triplet transitions feature significantly extended lifetimes and, like the TROSY experiment, can indicate a new approach for assignment and structural elucidation of proteins of increasing size by NMR.

For two *J*-coupled ^1^H nuclei, the nuclear singlet, a spin-exchange antisymmetric state, features slow-paced transitions to nuclear triplet states, which are spin-exchange symmetric. The new spectroscopic method to detect long-range interactions for proteins via their effect on LLC’s, exploits nuclear magnetic transitions similar to electronic phosphorescence. Transitions between nuclear singlet and triplet configurations were first observed as low-frequency oscillations in low magnetic fields (4). In high magnetic fields, these transitions offer extended time-frames for Overhauser-type transfer in the rotating frame (6–8) featuring, in high-resolution NMR magnets, decays up to 9 times slower than those of standard NMR transitions. LLC’s can be sustained with various methods in 2D experiments affording resolution at moderate amplitudes of the sustaining field (17). We report in this manuscript for the first time on the sensitivity of the sign of the detected signals for protein interactions to the reciprocal orientation of magnetic moments mixed in an LLC state. The stereospecific information for LLC interactions is obtained due to the fact that LLC’s are permutation-antisymmetric LLC’s. LLC’s are excited on naturally-occurring amino-acids in Lysozyme, a 14.3 kDa protein. The sign of these nuclear-permutation antisymmetric coherences can be switched by the experimenter and the interaction of LLC’s with different signs with surrounding hydrogens is a new way to establish the structural position of these atoms.

## Results and Discussion

We treat herein the interactions of protons with angular momenta ½*ħ* in the molecular framework of a protein. Pairs of protons Gly-H^α2,3^ in different glycine residues in Lysozyme are noted I and S, respectively. There are two possible orientations, (*α, β*)_*I,S*_, for each of their magnetic moments with respect to an external magnetic field, ***B***_0_. These two coupled spins that mainly interact with each other are described by the singlet-triplet wavefunctions (3), i.e., the nuclear spin-permutation antisymmetric singlet state, *S*_0_ = *N*(|*α*_*I*_*β*_*S*_ > − |*β*_*I*_*α*_*S*_ >), and the three symmetric triplet states, *T*_+1_ = | *α*_*I*_*α*_*S*_ >, *T*_0_ = *N*(|*α*_*I*_*β*_*S*_ > + | *β*_*I*_*α*_*S*_ >), *T*_−1_ = | *β*_*I*_ *β*_*S*_ >, with *N* = 2^−1/2^ (Figure 1). The decays of singlet-state based populations and transitions are the least perturbed by spin-permutation symmetric interactions, such as the dipole-dipole interaction between the two nuclei (3,4). Collective spin order with long lifetimes compared to classical spin order can be excited based on the population differences between singlet and triplet states, provided the external magnetic field is removed or eclipsed by strong radio-frequency irradiation (5).

**Figure 1.**
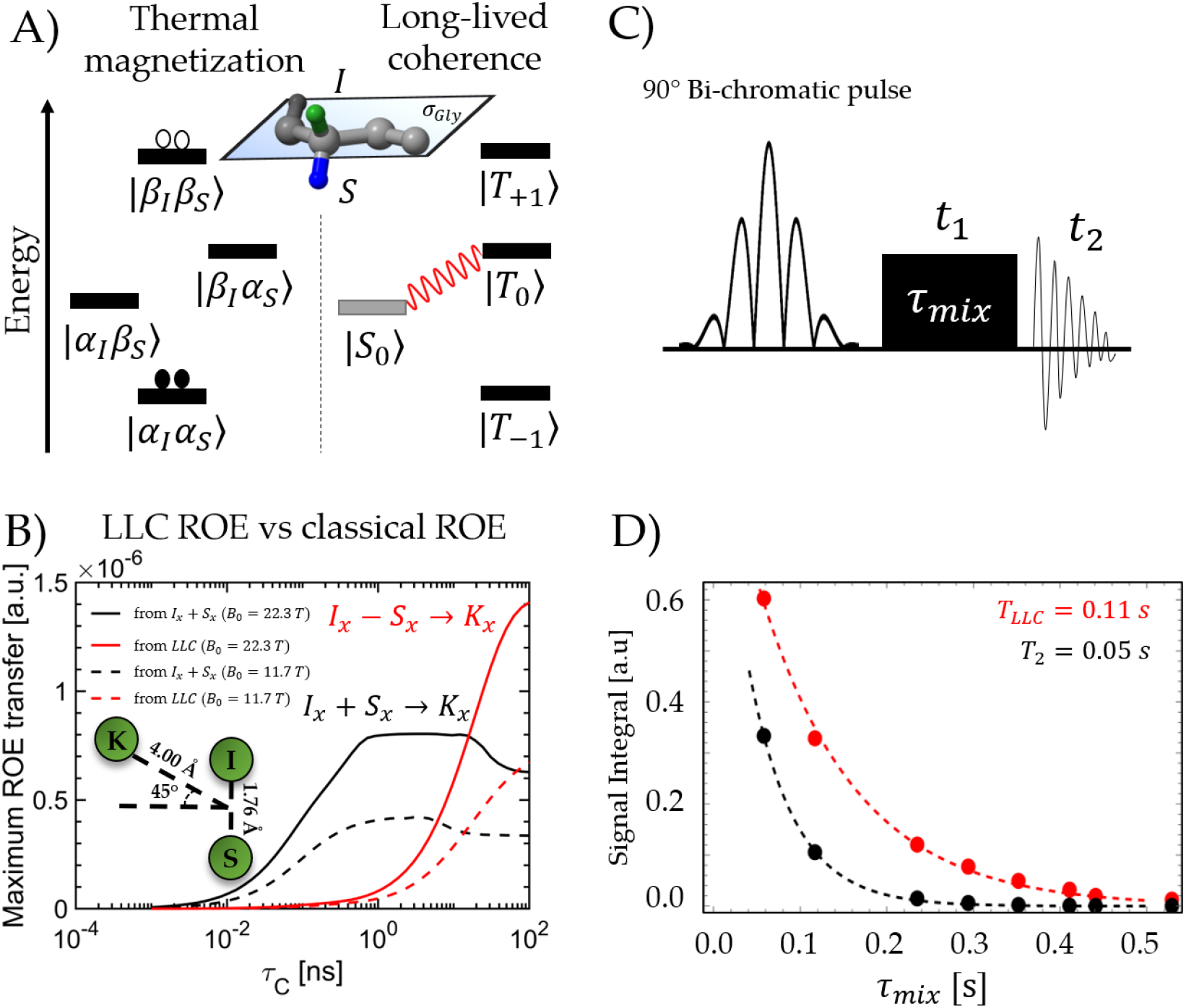
A) Long-lived coherences are quantum superpositions of singlet and triplet states of glycine *J*-coupled H^α 2,3^ protons. The energy level diagram of spin states of a *J*-coupled two-spin system in the weak-coupling and strong-coupling compared to the chemical shift difference between the two spins is shown. B) Theoretical dependence of LLC-based (red curve) and standard dipolar transfer (black curve) of transverse spin order from Gly aliphatic protons to a neighbour ^1^H spin, denoted K; Numerical simulations (*Spinach*^29^) describe the dependence of maximum rotating-frame Overhauser (ROE) transfer towards a third spin K, starting from 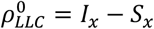 (red) compared to 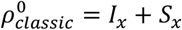 (black), as molecular tumbling slows down (the rotational correlation *Δ*_c_ time increases). The transfer between (I,S)-LLC and neighboring spin K gradually increases and surpasses the classical ROE transfer. Simulated geometrical parameters of the three spin systems are provided as an inset. C) The 1D/ 2D pulse sequence used to excite LLC’s on targeted Gly-H^α^ resonances. The initial phase offset between the two components of the 90° bi-chromatic pulse can be adjusted to generate either long-lived coherences (‘LLC’ experiment), or classical transverse coherence (“REF” experiment). The continuous irradiation sustain time is used as mixing time, *Δ*_mix_ (for 1D experiments) or indirect dimension evolution time, t_1_ (for 2D experiments). Other relevant parameters are: #*F*1 = 64; #*F*2 = 2048; *ns* = 32;. D) Signal decay of the LLC versus transverse magnetization during the spin-lock period (*τ*_*mix*_) measured at 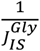 intervals.

### Long-lived coherences

The radio-frequency magnetic field sustaining condition described above equally enables the observation of coherences with relaxation time constants that are resilient to dipolar interactions, namely singlet-triplet transitions (Figure 1A), also known as long-lived coherences (LLC’s):

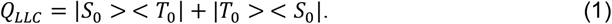

Expressed in terms of Cartesian operators (6,18), LLC’s have a real component parallel to the radio-frequency irradiation field, 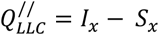 and one imaginary component with antiphase components composed of operators that are transverse to this field,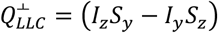.

The sensitivity of LLC’s to the presence of nearby nuclei yields new information compared to classical coherences. The interaction of an LLC superposition with a neighboring spin has been described both theoretically and via experiments for small-molecule systems (7). The dependence on the effective correlation time modulating the interaction, Δ_c_, of the transfer of magnetisation from LLC to a neighbour situated at a distance of 4 Å from the average position of Gly-H^α2,3^ is compared to the classical transfer of transverse magnetisation in Figure 1B. At a magnetic field corresponding to proton Larmor frequency of 950 MHz, proteins of the molecular weight of Lysozyme can be expected to feature enhancements of LLC-based transfer compared to classical transverse transfer.

Long-lived coherences can be accessed via guided magnetisation evolution (‘spin dynamics’) with user-applied radio-frequency pulses, starting from *I*_*z*_ + *S*_*z*_. This is performed using the 1D pulse sequence shown in Figure 1. LLC’s decay according to their auto-relaxation rate constants, *R*_LLC_ (Supporting Information, (7)). The Liouvillian of a 3-spin system (I = Gly-H^α2^, S = Gly-H^α3^, K = neighbor proton) shows that the frequency of oscillation is *v*_LLC_ = *J*_IS_ when *J*_IS_ is the dominant coupling for the spin system and *R*_LLC_ is the relaxation rate constant, which effectively describes the decay of the signal. We probed ROE transfer by targeting several glycine residues with the pulse sequence described in Figure 1C where we adapted the 90° selective pulse for each spin system, with an excitation profile described in Figure 1D. In the experimental methods, the LLC evolution time was chosen as multiples of the LLC period. This enabled us to derive the relaxation time constant from the fit (19,20) of LLC intensities as a function of time (Figure 1D). The relaxation time constant of LLC’s was found to be *T*_LLC_ = 110 ms, more than two times as long as the relaxation time constant of standard transverse magnetisation M_x_ = I_x_ + S_x_, locked along the x axis of the transverse plane, *T*_1ρ_ = 50 ms.

### Magnetisation transfer in 1D and 2D LLC-ROE build-ups

1D ROE spectroscopy is a fast way of probing interactions. The relaxation rate constants of LLC’s are driven by the dipolar interaction between coupled spins I and S, as well as by interactions with external relaxation sources, K_i_ (21). By starting from *I*_*x*_ − *S*_*x*_, the build up of polarization at the K resonance will change sign if K is closer to the S spin than to the I spin (See Supporting Information for angular dependence of LLC-ROE transfer). Signal intensity evolution during various mixing times *τ*_*mix*_ in LLC-ROE experiments starting from 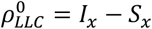 were compared with reference 1D ROE experiments starting from 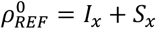. The results for G49, G67, G117 and provided in Figure 2. The sign of several cross-peaks changes in LLC-ROE with respect to reference ROE-1D - for example D119-H^α^, N46-H^α^, D66-H^α^, T51-Hγ^2^ in Figure 2 -due to the inversion of the selected glycine’s proton closest to the corresponding neighbouring proton. Furthermore, for several cross-peaks, e.g., G117-LLC/W111-H^ζ2^ in Figure 2, we observed an enhancement of the Overhauser transfer when starting from LLC compared to classical coherences, as predicted by numerical simulations and theory.

**Figure 2.**
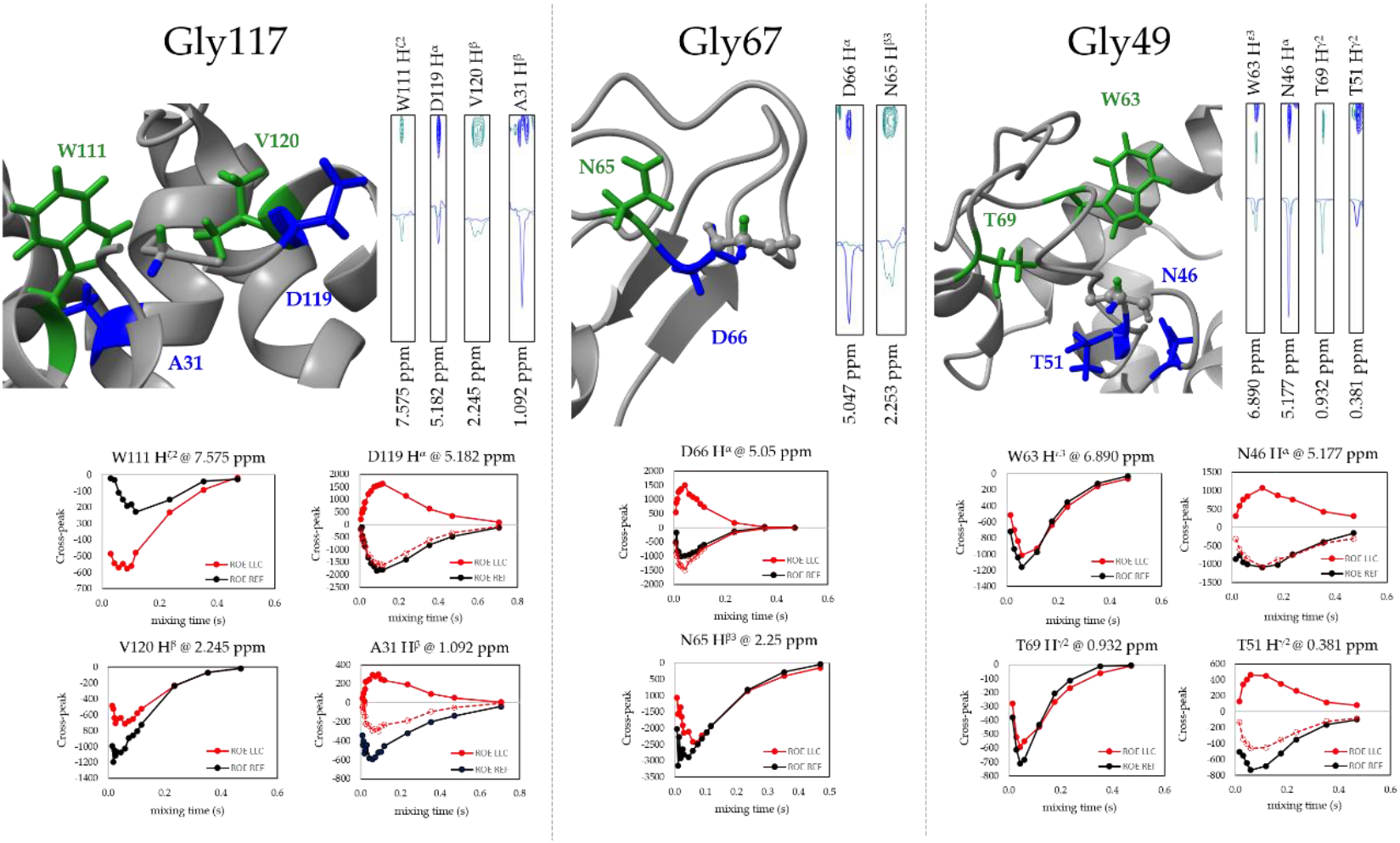
1D and 2D observation of Gly LLC-ROEs in Lysozyme for G117, G67, G49. ‘ROE LLC’ (red) versus ‘ROE reference’ (black) build-up experiments for G117 (left), G67 (middle) and G49 (right) with highlighted spatial neighbouring residues in the Lysozyme structure where Overhauser transfer is detected. The 1D signals were obtained based on LLC’s excited with the selective pulse shown in Figure 1C) both for the 1D version (build-ups) and 2D version (cross-signals). F_1_-stripes through the 2D ROE LLC experiments are superimposed with classical 2D ROESY slices at the frequency of aliphatic Gly-H^α2^ (green) and H^α3^ (blue). The change in sign of the cross-peak build up matches the change in relative intensity of the classical ROESY stripes. This confirms via the classical ROESY-2D experiment that atoms closer to H^α2^ have negative LLC build-ups, while the ones closer to H^α3^ have positive LLC build-ups. Detected ROE signals between G117 and residue D119-H^α^ (closer to H^α2^ than to H^α3^) are positive, therefore the plot of inverted intensities is also shown to facilitate comparison with classical build-up. The protein structure outlines Gly C^α^-H^α2^ bonds (green) and C^α^-H^α3^ bonds (blue), as well as interacting residues close in space to H^α2^ (e.g, W111 in green) respectively close in space to H^α3^ in blue (e.g., D119-H^α^).

Thus, dipolar-coupled spins can be mapped in the protein topology with respect to the symmetry plane of the targeted Gly methylene protons by the sign of their cross peak in an LLC experiment. The build-up of magnetisation originating on G117 encoded LLC’s at the frequency of a neighbouring spin K (Figure 2) has been probed as a function of the transfer time for neighbours W111-H^ζ2^, D119-H^α^, and others. Maximum transfers are found for build-up times of circa 100 ms. For the case of G117-H^α1,2^ transfer to W111-H^ζ2^, the intensity of the transfer from LLC-based configurations is two times that of the transfer based on standard transverse configurations, while for other neighbours D119-H^α^ the two types of transfer yield comparable results. A stronger intensity for LLC-based transfers is also found in the case of G67-H^α1,2^ transfer to D66-H^α^, as mentioned above. This is explained by the geometries of the (I,S,K) configurations in the two cases (Supporting Information). The transfer from Gly protons to neighbours is oscillating in the fast-diffusion limit and the oscillations are dampened by relaxation in the slow diffusion limit.(7) The intensity of observed transfers in Lysozyme is affected by the different components of the overall tumbling of the protein and local anisotropic motions. This sets effective local correlation times for each glycine interaction.

The magnetic interactions at the positions of Gly-H^α2^ and Gly-H^α3^ was also be probed using 2D spectroscopy. Signals are encoded by the sequence in Figure 1 for the Gly source protons as well as for the interacting partner protons at their respective frequencies in the direct dimension and at the frequency *v*^LLC^ = *J*_IS_ in the indirect dimension. Positive or negative intensities are expected according to the position of the interaction partner on one side or the other of the symmetry reflection plane of (H^α2^, H^α3^). Variations of the signal positions in the indirect dimension can occur as the oscillations are dampened by the relaxation of each individual spin (Figure 2).

A high number of ROE’s (11) could be observed with positive and negative signs, and the relative positions with respect to G49-H^α 2,3^ are confirmed by the relative intensities of the lines from classical 2D-ROESY spectroscopy at the frequencies of H^α2^ and H^α3^ in the indirect dimension. The majority of the detected LLC-ROE interaction signals were consistent with structural statistics in terms of angle and distance (Figures S2-S4 of Supporting Information). For instance, most of the 50 NMR structures in the available dataset were consistent with the stereospecific assignment of the interactions G49-LLC / N46-H^α^ (positive signal for a majority of positive φ angles in the 50 structures) and G49-LLC / T69-H^γ2^ (negative signal for a majority of negative φ angles in the structures), G67-LLC / N65-H^β3^ (positive signal, positive φ angles) and G67-LLC / D66-H^α^ (negative signal, negative φ angles). In cases when the LLC-interacting proton is situated very close to the symmetry plane, yielding very similar intensities in the two lines from classical ROESY-2D, the intensity of the 2D LLC-ROE is very small, which also yields valuable structure information.

In conclusion, we show that the lifetimes of ^1^H-based long-lived coherences in a 14.3 kDa protein can be used to transfer magnetisation to neighbor protons in the protein structure. Structural information involving distance and angle constraints between Gly aliphatic protons in loops and secondary structure elements can be obtained on this basis. Stereospecific assignment is obtained, as spin-permutation antisymmetric long-lived coherences yield signals with opposite signs for interactions with neighbours closer to H^α2^ and, respectively, H^α3^. The enhanced magnetisation transfer based on LLC-ROE compared to standard ROE in several residues may constitute the basis for new structural determination methods in large proteins where short relaxation lifetimes are a limiting factor. The method can be applied to obtain distance information for protein interactions in cells, as no isotopic enrichment is required - only hydrogen spins are used.

## Materials and Methods

The Lysozyme protein (29 mg, MW= 14300 g.mol^-1^) with natural-abundance spin isotopes was dissolved in D_2_O (0.55 ml). NMR spectra were recorded at T= 309 K on a Bruker Avance spectrometer operating at B0 = 22.325 T, i.e., at the Larmor proton frequency v_0_ = 950 MHz, equipped with a cryogenically-cooled probehead. Spectral intensities were extracted using TopSpin and Mestrenova. The dependences of spectral intensities on evolution delays were fitted using the dedicated Matlab function and errors were calculated from a Monte-Carlo analysis performed using 100 variations within the spectral noise level for each fit. Proton reference 1D spectra and LLC experiments were recorded with 8 transients and a recovery delay of 8 s. The (H^α1^, H^α2^) pairs in Gly feature ^2^*J*_IS_ coupling values *J*_IS_ = 17 Hz and frequency differences of Δ_*V* IS_ at the given B_0_ value provided in the supporting material. To record an LLC’s experiment with the inversion of one Gly-H^α^, we used the pulse sequence describes in Figure 1. The carrier frequency of the excitation pulse was placed on the most downfield proton and the continuous wave irradiation pulse was set in the middle of the two Gly-H^α^ doublets. The excitation range of the selective pulse was +/-70 Hz for each Gly proton, simultaneous excitation of I_x_ and -S_x_ further ensuring source selection. Of the twelve Gly residues in Lysosyme, spectral dispersion was sufficient to excite the residues discussed in the text. In practice, LLC terms are excited by continuous-wave irradiation with an amplitude γ(^1^H)***B***_1_/(2π)= 4 kHz. The duration of the applied radio-frequency sustaining was varied to match the desired mixing time. The spin dynamics behavior was simulated using SpinDynamica, ‘Spinach’ Matlab libraries and GAMMA C++ libraries (18,22,23) within a three-spin system (I,S,K) similar to the Gly-H^α^1, Gly-H^α2^, and neighboring (K) spin system.

## Acknowledgments

We are grateful to Massimo Lucci for assistance with the experiments, to Isabella Felli and Lucia Banci for disccussions, to Marco Allegrozzi, Rebecca del Conte for assisting with sample and manuscript data preparation. We acknowledge the iNEXT-Discovery (H2020, contract No 871037, Project ID-18523) hosted at the Magnetic Resonance Centre (CERM), University of Florence, for NMR time and funding from the Romanian Ministry of Research, UEFISCDI PN-III-P4-ID-PCE-2020-2642, UEFISCDI PN-III-P4-PED-565, Extreme Light Infrastructure Nuclear Physics (ELI-NP) Phase II, a project co-financed by the Romanian Government and the European Union through the European Regional Development Fund and the Competitiveness Operational Programme (1/07.07.2016, ID 1334) Romanian Ministry of Research, Equipments of National Interest (IOSIN).

