## Supporting_info for "Long-lived coherences for magnetic interactions in proteins"

for

Table of Contents

1. Experimental workflow for selective long-lived coherences experiments in large proteins
2. Theoretical analysis of rotating frame Overhauser transfer from long-lived coherences
3. Angular dependence of ROE LLC versus classical ROE transfer
4. Spatial distribution of neighboring protons around investigated glycine residues
5. 2D ROE LLC experiments

1. NMR experimental workflow for selective long-lived coherence experiments in large proteins

The workflow employed for the identification of glycine resonances and selective LLC excitation of glycine residues in Lysozyme at 950 MHz.

Step 1. Broadband excitation and detection of LLS. Glycine resonances are identified (blue) in this way, but not clearly assigned.

Step 2. Based on knowledge of resonance frequencies acquired at Step 1, selective excitation and detection of Gly49 LLS signal (green) utilizing the frequency-selective pulse sequence. This method allows for unambiguous assignment of coupled spins resonances in each glycine using an LLS filter.

Step 3. Once both resonances of one glycine residue are assigned, a 50 ms bichromatic EBURP2 pulse (with initial frequency offset) allows simultaneous excitation of transverse magnetization in two spectral regions with opposite sign, leading to LLC excitation of the selected Glycine residue (red). The excitation spectral regions (with a spectral width around 70 Hz each) are centered on the previously assigned glycine resonances.

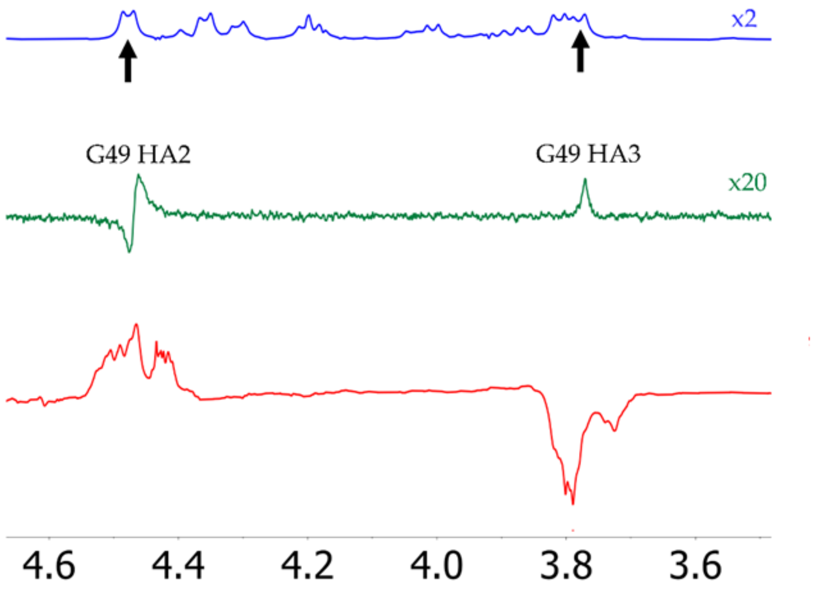

Figure 1S. (top) Protein Gly signals excited by LLS (blue); (middle) selective excitation and detection of Gly49 LLS signal (green) using the frequency-selective pulse sequence; (botton) bi-chromatic excitation of LLC for Gly-49.

Table S1. Assigned glycine resonances using the workflow method described above

| Residue | HA2 (ppm) | HA3 (ppm) | Δν (Hz) @ 950 MHz |
|---|---|---|---|
| G4 | 4.384 | 4.034 | 332.5 |
| G16 | 4.177 | 3.971 | 195.7 |
| G49 | 4.471 | 3.773 | 663.1 |
| G67 | 4.193 | 3.887 | 290.7 |
| G117 | 4.185 | 3.860 | 308.8 |
| G126 | 4.351 | 3.806 | 517.8 |

#### 2. Theoretical analysis of rotating frame Overhauser transfer from long-lived coherence

A complete theoretical overview is given in reference 7 of the main text. In the following we consider evolution of a system of three spins, *I*, *S*, and *K*, with relaxation due to dipolar interaction, and *J*-coupling between *I* and *S*. To simplify notation, we introduce for the expectation values of coherences involved in the ROE process the following symbols: $K = \langle K_x \rangle$, $I = \langle I_x \rangle$, $S = \langle S_x \rangle$, $Q_+ = \langle I_x + S_x \rangle$, $Q_{LLC}^{x} = \langle I_x - S_x \rangle$, and $Q_{LLC}^{yz} = \langle 2I_yS_z - 2I_zS_y \rangle$. Relaxation in the *I*-*S*-*K* system is described by auto-relaxation rates $\rho_I = \rho_S, \rho_K$ and cross-relaxation rates $\sigma_{IS}, \sigma_{IK}, \sigma_{SK}$. Neglecting coherent evolution due to the *J*-coupling between the *I* and *S* spins, it is found that

$$dI/dt = -\rho_I I - \sigma_{IS} S - \sigma_{IK} K, \qquad [1]$$

$$dS/dt = -\rho_S S - \sigma_{IS} I - \sigma_{IK} K, \qquad [2]$$

From Eq. [1,2], the evolution of $Q_{LLC}^{x}$ is given by

$$dQ_{LLC}^{x}/dt = -\rho_{LLC} Q_{LLC}^{x} - \sigma_{LLC}^{K} K \qquad [3]$$

where $\rho_{LLC} = \rho_I - \sigma_{IS}$ and $\sigma_{LLC}^{K} = \sigma_{IK} - \sigma_{SK}$. The equation for the change rate of $K$,

$$dK/dt = -\rho_K K - \sigma_{IK} I - \sigma_{SK} S, \qquad [4]$$

can be recast as

$$dK/dt = -\rho_K K - \frac{1}{2}\sigma_{LLC}^{K} Q_{LLC}^{x} - \frac{1}{2}\sigma_+ Q_+, \qquad [5]$$

where $\sigma_+ = \sigma_{IK} + \sigma_{SK}$. Because of the appearance of $Q_+$ in Eq. [7], an additional equation must be added which, according to Eq. [1] and [2], is

$$dQ_+/dt = -\rho_+ Q_+ - \sigma_+ K, \qquad [6]$$

where $\rho_+ = \rho_I + \sigma_{IS}$. However, according to the experimental procedure for producing LLC, initially $Q_+(0) = \langle I_x + S_x \rangle(0) = 0$ and it can be expected that $Q_+(t)$ remains small throughout the ROESY irradiation. Therefore, we can ignore Eq. [6] and neglect $\sigma_+ Q_+$ term in Eq. [7], such that we are left with the system of equations

$$dQ_{LLC}^{x}/dt = -\rho_{LLC}Q_{LLC}^{x} - \sigma_{LLC}^{K}K, \qquad [7]$$

$$dK/dt = -\rho_K K - \frac{1}{2}\sigma_{LLC}^{K}Q_{LLC}^{x}. \qquad [8]$$

This system of equations can be solved and the ROE transfer to *K*, starting from the initial condition $Q_{LLC}^{x}(0)$=1, $K(0) = 0$ is described by

$$K(t) = \frac{\sigma_{LLC}^{K}}{2A}e^{-(\rho_{LLC}+\rho_K)t}\left[e^{-At/2} - e^{+At/2}\right] \qquad [9]$$

where $A = \sqrt{(\rho_K - \rho_{LLC})^2 + 2\sigma_{LLC}^{K}}$.

In comparison, if one prepares the system as $Q_+(0) = 1, K(0) = 1$, it can be expected that $Q_{LLC}^{x}(t)$ remains small throughout the ROESY irradiation. Therefore, we can ignore Eq. [6] and neglect $\sigma_+Q_+$ term in Eq. [3], such that we are left with the system of equations

$$dQ_+/dt = -\rho_+Q_+ - \sigma_+K$$

$$dK/dt = -\rho_K K - \frac{1}{2}\sigma_+Q_+.$$

The ROE transfer to *K*, starting from the initial condition $Q_+(0) = 1$, $K(0) = 0$ is given by

$$K(t) = \frac{\sigma_+}{2B}e^{-(\rho_+ +\rho_K)t}\left[e^{-Bt/2} - e^{+Bt/2}\right] \qquad [10]$$

where $B = \sqrt{(\rho_K - \rho_+)^2 + 2\sigma_+}$.

#### 3. Angular dependence of ROE LLC versus classical ROE transfer

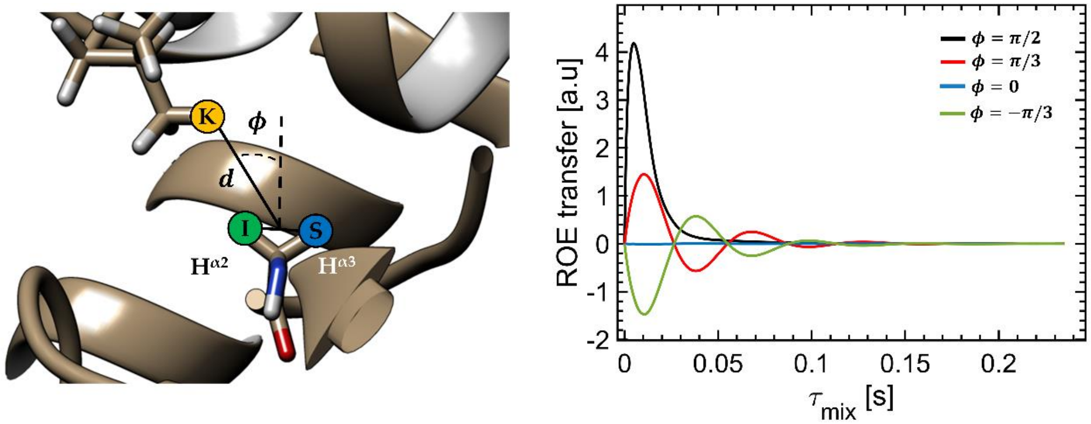


Fig S5. Angular dependence of ratio $(f)$ between maximum magnetization transfer from ROE LLC and classical ROE as function of $\phi$ angle at $\tau_C = 15\ ns$ rotational correlation time. The $\{I, S, K\}$ spin system has the following geometrical constrains: the internuclear distance between I and S spins is 1.76 Å and the distance between K spin and the middle point of the I-S segment is 2.3 Å.

### 4. Spatial distribution of neighboring protons around investigated glycine residues

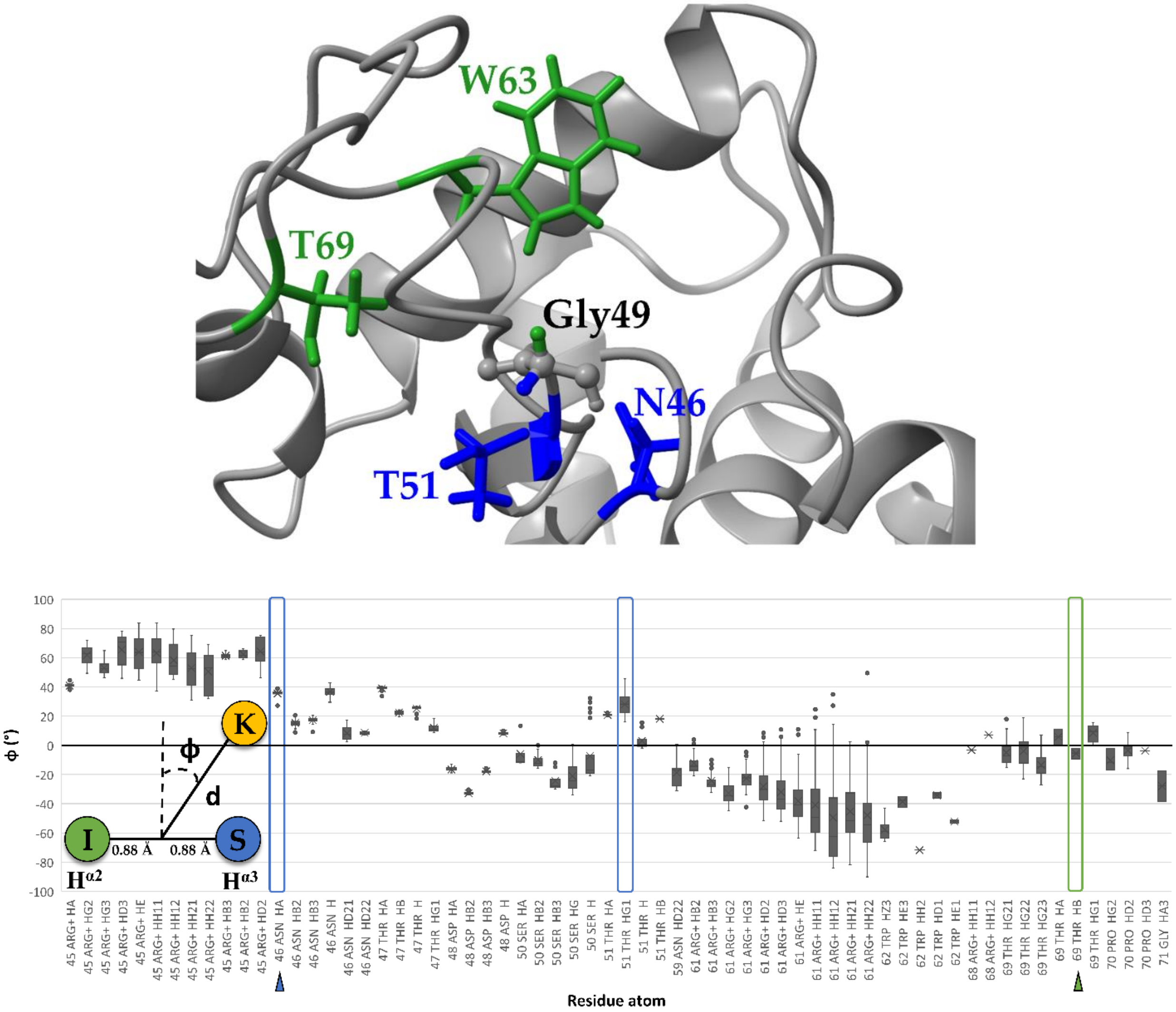


FigS2. Angle distribution of hydrogen atoms within an 8 Å radius from Gly49 residue over the 50 structures of Lysozyme from the "1E8L" PDB file. The "median" refers to the distance between the neighboring atom and the middle point of the Gly-HA2/HA3 internuclear segment. The $\phi$ angle is the angle between the median and the perpendicular bisector of the Gly-HA2/HA3 internuclear segment.

The spatial distribution of three neighboring protons with respect to the Glycine 49 (Figure S2) is confirmed by the sign of the LLC ROE transfer (Figure 2 in the main text). The spins closer to the Gly-HA3 (S spin in blue) display a negative ROE magnetization transfer. This is the case for Asn46 HA and Thr51 HG1. For Thr69 HB, the ROE build up has a different sign in agreement with the closer proximity of this spin to Gly-HA2 (I spin in green). The angular dependence of the ROE sign as described above is confirmed for the Gly67 residue (Figure S3).

### 5. 2D ROE LLC experiments

Gly49 ROE LLC 2D

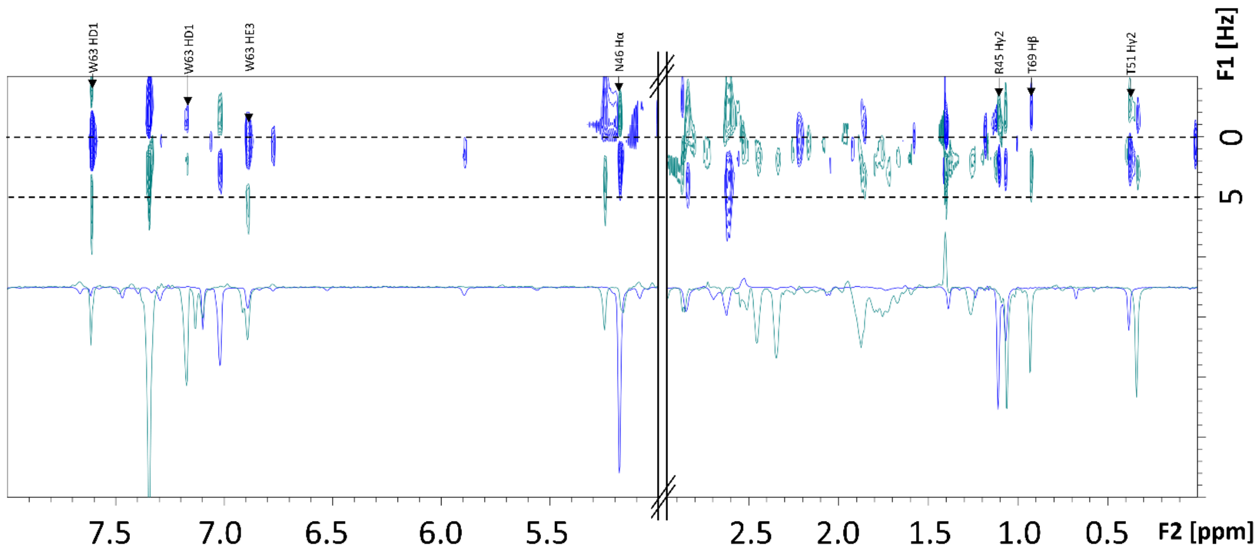


Fig S6. ROE LLC 2D spectrum (recorded with pulse sequence from Fig 1C) superimposed with slices from ROESY 2D spectrum for Gly49 residue. Spatial neighbors within a 10 A radius were assigned.

Gly117 ROE LLC 2D

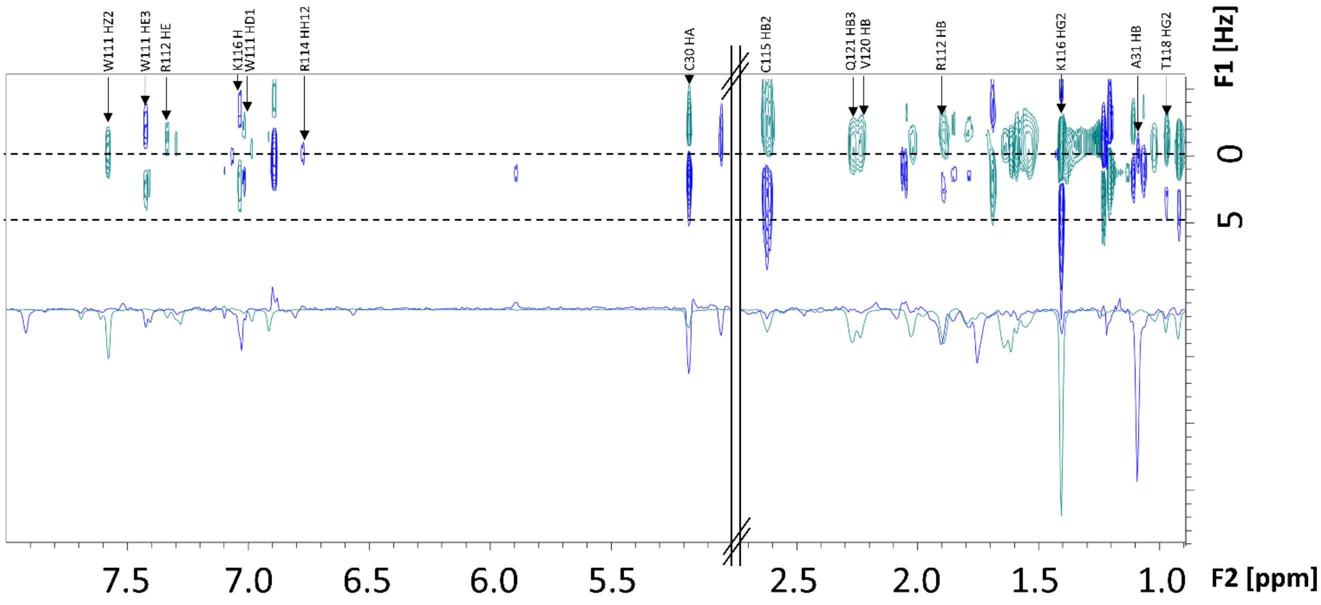


Fig S7. ROE LLC 2D spectrum (recorded with pulse sequence from Fig 1C) superimposed with slices from ROESY 2D spectrum for Gly117 residue. Spatial neighbors within a 10 A radius have been assigned.
